# A conserved root-knot nematode effector targets plant kinesin light chain related proteins to promote parasitism

**DOI:** 10.64898/2026.03.02.708937

**Authors:** Salomé Soulé, Sprine Misiani, Karine Mulet, Isabelle Mila, Jess Philippon, Claire Caravel, Joffrey Mejias, Stéphanie Jaubert, Pierre Abad, Nemo Peeters, Bruno Favery, Michaël Quentin

## Abstract

Root-knot nematodes (RKNs) are obligate plant parasites that establish biotrophic interactions with a wide range of host plants, causing major crop losses worldwide. During infection, RKNs secrete effector proteins from their esophageal glands into root tissues *via* a stylet. These effectors target specific subcellular compartments and manipulate host cell functions to induce the formation of hypertrophied and multinucleated giant cells, which serve as feeding sites essential for nematode development. Here we describe a new RKN-specific effector, EFFECTOR17 (EFF17), conserved in the five most damaging RKN species: *Meloidogyne incognita, M. enterolobii*, *M. arenaria*, *M. javanica*, and *M. hapla*. *In situ* hybridization showed that *EFF17* genes are specifically expressed in the esophageal glands of various RKN species. Silencing *EFF17* in *M. incognita* reduced its reproduction on *Nicotiana benthamiana*, demonstrating a key role in parasitism. Yeast two-hybrid and split luciferase assays revealed that tomato and *Arabidopsis* KINESIN LIGHT CHAIN RELATED (KLCR)/CELLULOSE-MICROUBULE UNCOUPLING (CMU) proteins are host targets of MiEFF17. Arabidopsis *klcr* mutants developed significantly fewer galls and egg masses. Our findings propose that EFF17 manipulates KLCR function to promote parasitism.

## Introduction

Plant-parasitic nematodes constitute a major threat to global agriculture, collectively causing billions of dollars in crop losses annually (Jones *et al*., 2013). Among them, root-knot nematodes (RKNs) of the genus *Meloidogyne* are the most economically damaging and globally distributed, an have the ability to infect thousands of plant species (Jones *et al*., 2013). The most destructive RKN species include the tropical species *Meloidogyne incognita*, *M. arenaria* and *M. javanica*, as well as the temperate species *M. hapla.* In addition, emerging species such as *M. enterolobii*, are particularly problematic in intensive cropping systems (Blok *et al*., 2008; Castagnone-Sereno *et al*., 2013; Jones *et al*., 2013).

RKNs are obligate sedentary endoparasites that establish long-term feeding relationships within roots (Favery *et al*., 2020). Following penetration and migration toward the vascular cylinder, infective second-stage juveniles induce the neoformation of specialized feeding cells called giant cells. These hypertrophied, multinucleate, and metabolically hyperactive cells serve as nutrient sources supporting nematode development and reproduction. Concurrently to giant cells development, surrounding cells proliferate, vascular tissues are extensively reorganized, and xylem and phloem elements expand (Bartlem *et al*., 2014). Hyperplasia of root tissues leads to the formation of a characteristic gall, a distinctive feature of *Meloidogyne* infection, which impairs water and nutrient uptake and consequently reduce plant growth. Within the gall, the nematode develops through successive molts before reaching the adult reproductive female stage that will release eggs in a gelatinous matrix at the root surface. Despite extensive research, the molecular determinants from both the nematode and the host that are required for the establishment and maintenance of parasitism remain poorly understood (Bali and Gleason, 2024).

The ontogenesis of giant cells requires extensive transcriptional, metabolic, and structural reprogramming of host root cells (Cabrera *et al*., 2014; Favery *et al*., 2016; Olmo *et al*., 2020). Hallmarks of this developmental conversion are dramatic changes in the organization of the plant cell wall, cytoskeleton and endomembrane system. Giant cells undergo a tremendous isotropic expansion accompanied by cell wall thickening and profound reorganization of microtubules (MTs) and actin filaments (Rodiuc *et al*., 2014; de Almeida Engler *et al*., 2004; de Almeida Engler *et al*., 2010). Pharmacological studies demonstrated that stabilization of MTs interferes with feeding site development, whereas controlled cytoskeletal destabilization can be compatible with nematode development, highlighting the importance of cytoskeletal dynamics during infection (de Almeida Engler *et al*., 2010). Similarly, genetic mutations affecting MT organization can impair feeding site development (Caillaud *et al*., 2008; Banora *et al*., 2011). Together, these findings provide compelling evidence that precise modulation of host cytoskeletal architecture is a critical target of RKN manipulation to ensure successful feeding site formation.

The initiation and maintenance of giant cells, unique to RKN parasitism, is achieved through the secretion into the plant of effector proteins mainly produced in esophageal glands, including one dorsal and two subventral glands, through a syringe-like stylet. Over the past decade, genomic and transcriptomic analyses have revealed an extraordinarily diverse effector repertoire in *Meloidogyne* spp., including proteins targeting hormone signaling pathways, RNA metabolism, protein turnover, redox homeostasis, or cytoskeletal organization (Vieira & Gleason, 2019; Rutter *et al*., 2022; Bali and Gleason 2024).

The plant cytoskeleton has emerged as a recurrent target of plant pathogen effectors (Park *et al*., 2018; Wang *et al*., 2022). Effectors from the sedentary endoparasite cyst nematodes and RKNs have been shown to interfere directly with actin and MT dynamics (Leelarasamee *et al*., 2018; Mei *et al*., 2018). Among cytoskeletal regulators, plant kinesin light chain-related proteins (KLCRs), also known as cellulose–MT uncoupling (CMU) proteins, play a critical role in stabilizing cortical MTs against lateral displacement driven by cellulose synthase complexes (Liu *et al*., 2016; Ganguly *et al*., 2020). In *Arabidopsis thaliana*, KLCR/CMU proteins contribute to proper cellulose deposition, MT stability, and cell morphogenesis (Liu *et al*., 2016; Ganguly *et al*., 2020; Yang *et al*., 2022). These proteins function as scaffolding components within multiprotein complexes that link MTs, actin filaments, and membrane contact sites, particularly at endoplasmic reticulum–plasma membrane interfaces (Zang *et al*., 2021; Wang *et al*., 2023). Noteworthy, KLCR proteins have been identified as hubs targeted by effectors from diverse microbial pathogens, including bacteria and oomycetes (Mukhtar *et al*., 2011; Weßling *et al*., 2014) and the *M. incognita* Minc00344 and *M. javanica* Mj-NULG1a effectors (Godinho Mendes *et al*., 2021), reinforcing the concept that cytoskeletal scaffolds constitute targetted regulatory nodes. Given the extensive ER proliferation and cytoskeletal reorganization observed in giant cells (de Almeida Engler *et al*., 2004; Miyashita & Koga, 2017; Orange *et al*., 2025), KLCR proteins represent interesting candidate susceptibility factors during RKN infection.

In the present study, we described a previously unannotated RKN-specific effector, EFF17, expressed in the subventral esophageal glands of the major *Meloidogyne* spp. We investigated the function of this pioneering effector by identifying its host targets using a yeast two-hybrid (Y2H) strategy. We established that *M. incognita* EFF17 (MiEFF17) interacts *in planta* with tomato and *Arabidopsis* KLCRs. The characterization of *Arabidopsis* mutants demonstrates the role played by KLCRs in susceptibility to *M. incognita*. Together, our findings identify EFF17 as a conserved *Meloidogyne* effector that targets cytoskeletal regulatory proteins central to cellulose–MT coordination allowing feeding sites development and supports the idea that diverse pathogens converge onto shared plant regulatory hubs to promote parasitism.

## Results

### EFF17 is an RKN-specific effector required for parasitism

Three genes are predicted to encode putative MiEFF17s in the *M. incognita* genome (Blanc-Mathieu et al., 2017): *Minc3s01763g26124/Minc17612/MiEFF17a*, *Minc3s00976g19489/MiEFF17b* and *Minc3s04571g36568/MiEFF17c*. MiEFF17a, MiEFF17b and MiEFF17c are 131-, 131- and 134-aa proteins, respectively and have an 18-aa or 20-aa signal peptide (SP) for secretion according to the PSORTII prediction tool (Horton and Nakai, 1997) (Fig. 1A; Supplementary Fig. S1). Mature EFF17 proteins share 87.3% amino acid (aa) identity a 45-48 aa C-terminal region enriched in glycine (G; 60%) and aromatic residues (F+W+Y; 31%) (Fig. 1A). The *EFF17* genes are specific to RKNs and could be recovered in every available RKN genome (Fig. 1B and 1C; Supplementary Fig. S1). Our previous RNAseq data indicate that all three *MiEFF17* genes, particularly *MiEFF17a* and *MiEFF17b*, are more strongly expressed during the parasitic stages (Fig. 2A; Blanc-Mathieu *et al*., 2017). *In situ* hybridization experiments were performed using a *MiEFF17a*-specific antisense probe showing that *EFF17* transcripts are produced specifically in the subventral glands of pre-parasitic J2 larvae *in M. incognita*, *M. enterolobii, M. arenaria* and *M. javanica* (Fig. 2B). No signal was detected in pre-J2s when using sense probes (Fig. 2B).

**Fig. 1.**
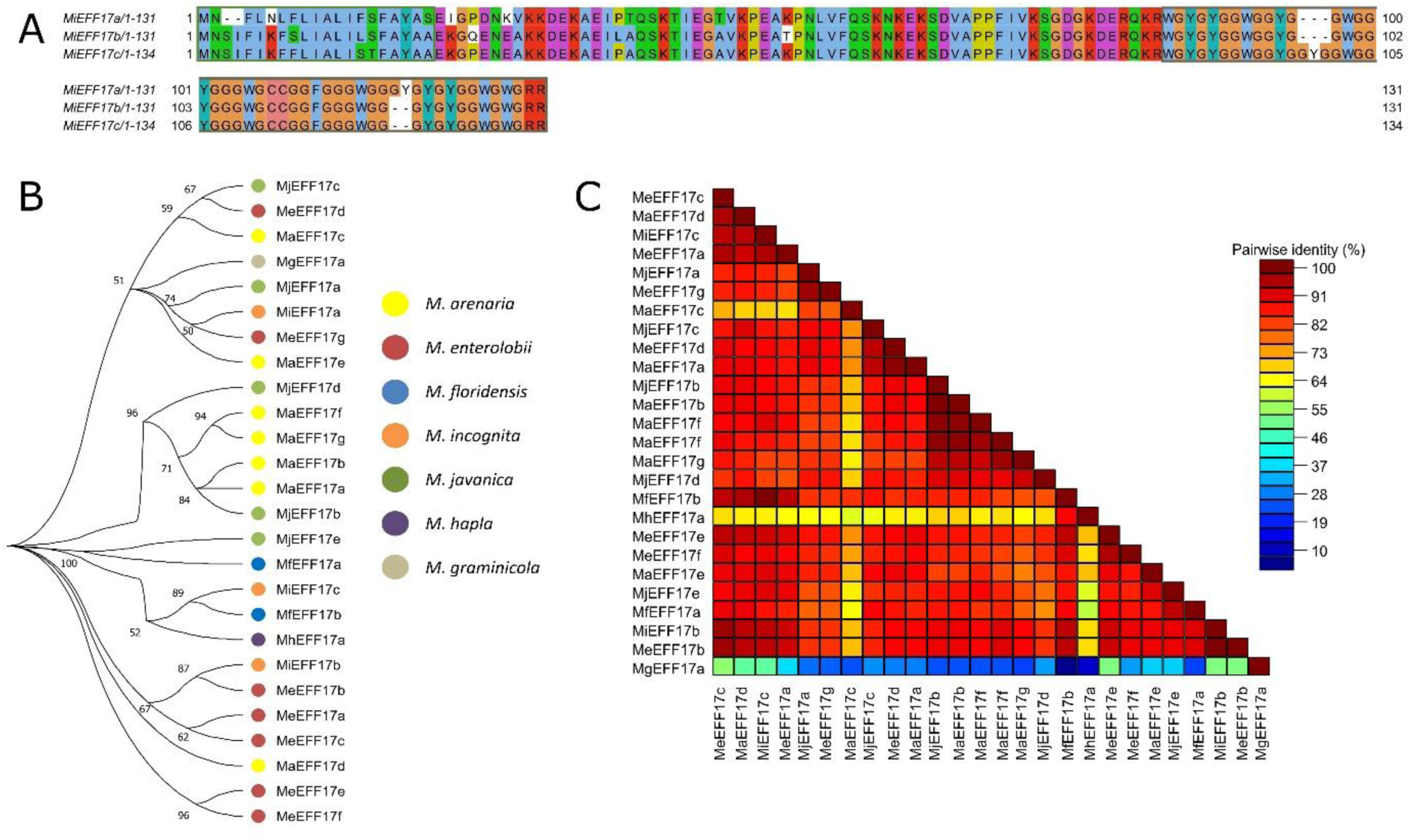
EFF17 is a pioneer conserved *Meloidogyne*-specific effector. **A,** Alignment of the MiEFF17 protein sequences. The green box indicates the position of the signal peptide for secretion. The brown box shows the C-terminus glycine-rich region. **B,** Phylogenetic tree of *Meloidogyne* spp. EFF17 amino-acid sequences. The percentages displayed next to each branch represent the number of tree replicates in which the associated taxa were assembled together in 1,500 bootstraps. The lengths of the branches are not proportional to phylogenetic distance. **C,** Pairwise amino acids sequence identity matrix for RKN EFF17 proteins.

**Fig. 2.**
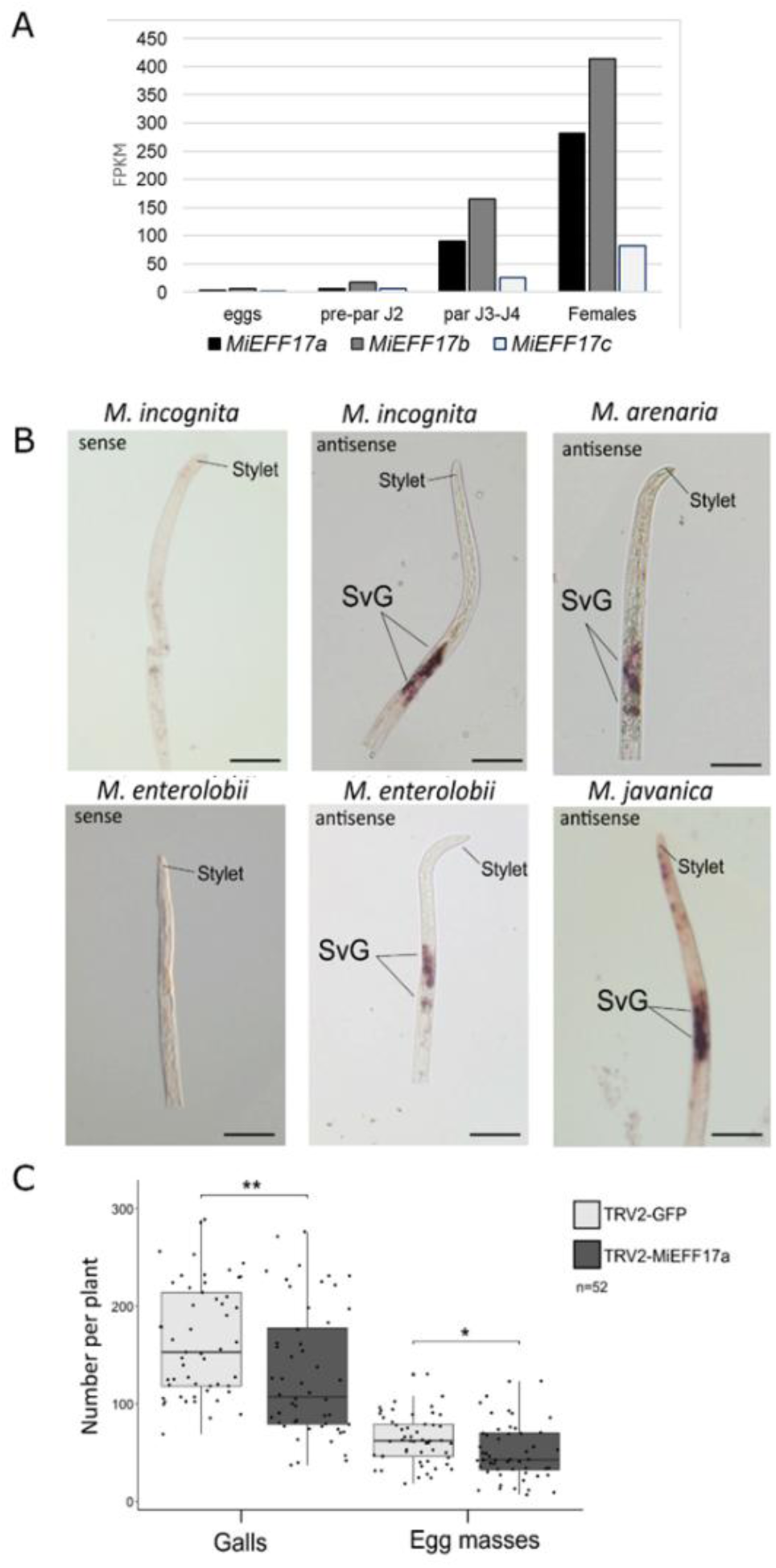
MiEFF17 is a secreted effector contributing to M. incognita parasitism. **A,** Transcript abundance of *MiEFF17a*, *MiEFF17b* and *MiEFF17c* in *M. incognita* eggs, pre-parasitic juveniles (pre-par J2), parasitic juveniles (par J3-J4) and females was retrieved from RNAseq data (Blanc-Mathieu et al., 2017) and is displayed as Fragments Per Kilobase per Million mapped fragments (FPKM). **B,** *In situ* hybridization (ISH) of digoxigenin-labelled *EFF17* probes in RKN second-stage juveniles (J2s). *EFF17* sense probe was used as a negative control giving no signal in *M. incognita* or *M. enterolobii* pre-parasitic J2. Using the antisense probe, *EFF17* transcripts specifically detected in the subventral esophageal glands (SvG) of *M. incognita*, *M. arenaria*, *M. enterolobii* and *M. javanica*. Bars, 20 µm. C, Infection test on *N. benthamiana* control plants (TRV2-GFP) and plants producing siRNA for the silencing of *MiEFF17* genes in *M. incognita* (TRV2-MiEFF17a). Boxplots represent the distribution of galls and egg masses six weeks after inoculation with 200 second-stage juveniles per plant across three biological replicates (n=52 plants). Box indicates interquartile range (25th to the 75th percentile). The central line within the box represents the median. Whiskers indicate the minimum and maximum values for the normal values present in the dataset. Statistical significance was assessed using non-parametric Mann-Whitney tests (*p<0.05; **p<0.01).

To demonstrate the role played by MiEFF17 during parasitism, we used a host-induced gene silencing (HIGS) approach. We silenced *MiEFF17* during *M. incognita* feeding on *Nicotiana benthamiana* inoculated with recombinant tobacco rattle virus (TRV) triggering virus-induced gene silencing (VIGS). A construct targeting the green fluorescent protein (GFP) transcript was used as a negative control. Six weeks post-inoculation with *M. incognita*, roots of *N. benthamiana* treated with a TRV meant to silence *MiEFF17* e exhibited a significantly decreased numbers of galls and egg masses when compared to the TRV2:GFP control (Fig. 2C). Together, these results indicate that *EFF17* genes encode RKN-specific effectors that are produced in secretory organs and are required for successful parasitism.

### MiEFF17 interacts with kinesin light chain-related proteins

To investigate the function of EFF17 *in planta*, we searched for its host targets using a yeast two-hybrid screen. MiEFF17a without its signal peptide was used as a bait to screen a cDNA library from healthy and *M. incognita*-infected tomato roots (Hybrigenics Service, France). In total, 91.8 million interactions between MiEFF17a and proteins encoded by the tomato cDNA library were screened. We recovered 29 clones capable of growing on selective medium lacking histidine. Nine clones were isolated carrying cDNAs corresponding to the three *Solanum lycopersicum KLCR* genes (Fig. 2A; Supplementary Fig. S2) named *SlKLCR1* (*Solyc08g008460*), *SlKLCR2* (*Solyc02g086840*) and *SlKLCR3* (*Solyc03g114320*). The imidazoleglycerol-phosphate dehydratase, captured three times here, was identified as the target of different effectors under our screening conditions and was considered as a false positive (Zhao *et al*., 2020; Mejias *et al*., 2021). The remaining candidate targets were recovered only once or twice. KLCRs were thus considered as the most likely host target of MiEFF17a.

The three *SlKLCRs* are diversely expressed during plant development (Supplementary Fig. S2-S4) and encode proteins owing characteristic tetratricopeptide repeat (TRP) domains. Similarly, AtKLCR1 (AT4G10840), AtKLCR2 (AT3G27960) and AtKLCR3 (AT1G27500) are encoded by three genes in *Arabidopsis* (Supplementary Fig. S5). To validate further the interactions between KLCRs and MiEFF17a *in planta*, we performed split luciferase assays. An interaction between two fusion proteins transiently expressed, *e.g.* RipG7 and its plant target MSKA (used as positive controls, Wang *et al*., 2016) fused with the N-terminal half (-Nluc) and the C-terminal half (-Cluc) of the luciferase, respectively, reconstitutes a functional enzyme, emitting light in the presence of luciferin that can be quantified with a luminometer. We transiently expressed in *N. benthamiana* leaf cells MiEFF17a-Nluc and Cluc-KLCRs fusion proteins and confirmed their integrity by performing western-blot analysis (Supplementary Fig. S6). This split-luciferase assay showed that co-expression of Cluc-KLCRs with MiEFF17a-Nluc produces a bioluminescence signal significantly higher than the Cluc-KLCRs/RipG7-Nluc combination, used as negative control. By contrast, as a positive control, RipG7 and its plant target MSKA fused with Nluc and Cluc, respectively, interacted when co-expressed *in planta* (Wang *et al*., 2016). AtKLCR1 and AtKLCR3 showed the strongest interactions with MiEFF17a-Nluc (Fig. 3B).

**Fig. 3.**
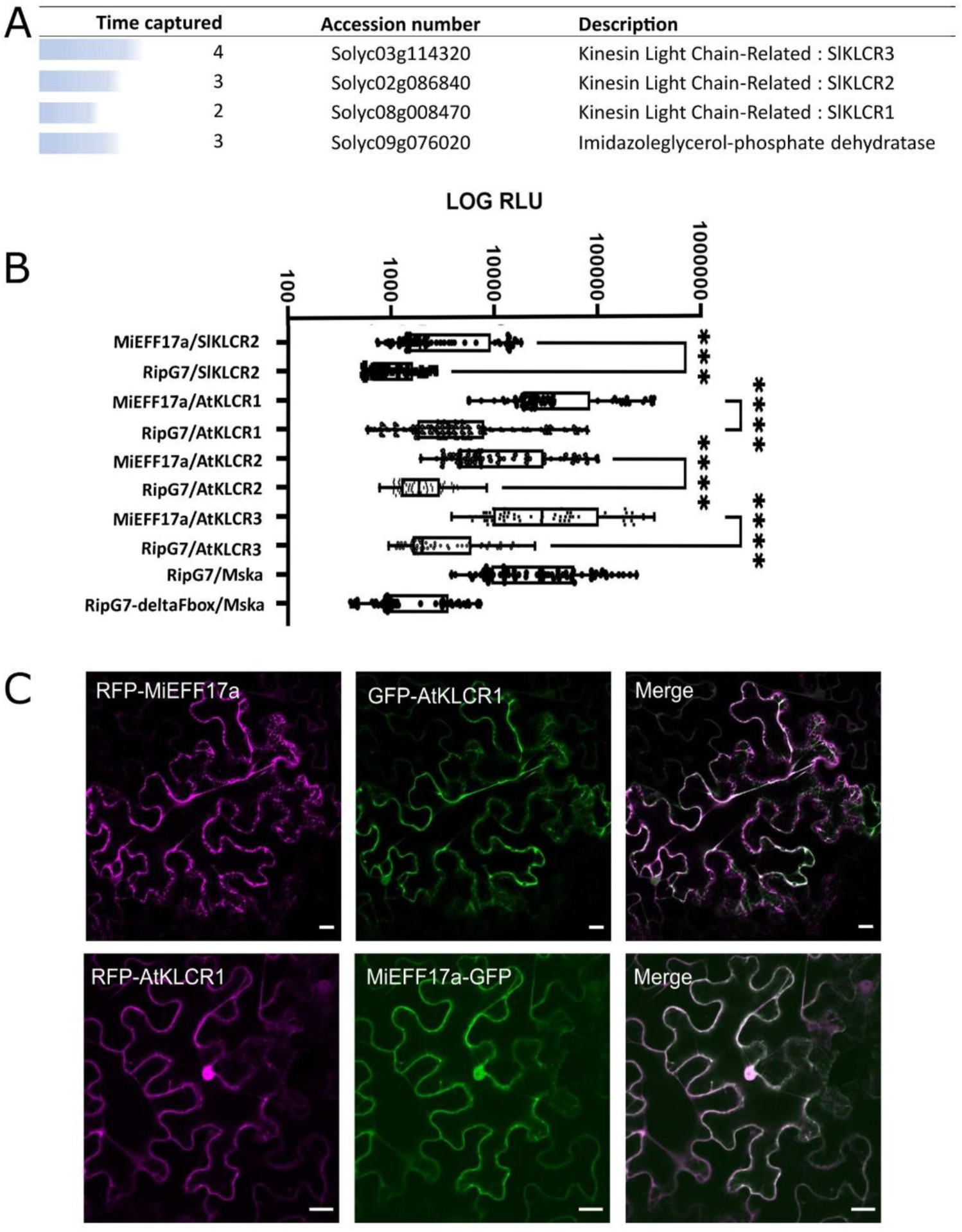
MiEFF17a physically interacts with host KLCRs. **A,** List of putative MiEFF17-interacting tomato proteins identified in our Y2H screen. **B,** A split luciferase assay confirmed the interaction between MiEFF17a and tomato and *Arabidopsis* KLCR. Cluc-KLCRs and MiEFF17a-Nluc were transiently co-expressed in *N. benthamiana* leaf cells and the light emitted by the reconstitution of the luciferase was quantified using a luminometer (relative light unit, RLU). The co-expression, of RipG7-deltaFbox-Nluc with all different Cluc-fusions was used as negative controls, while the co-expression of RipG7-Nluc with Cluc-MSKa was used as positive control. Experiment was conducted in three independent biological replicates, each containing 16 technical replicates. All data points are represented in this figure. Non-parametric Kruskal-Wallis statistical test was used (*** p-value < 0,001; **** p-value < 0,0001). **C,** Co-localization analysis of MiEFF17a and AtKLCR1. mRFP-MiEFF17a and eGFP-AtKLCR1 or MiEFF17-eGFP and mRFP-AtKLCR1 were transiently co-expressed using *Agrobacterium* in *N. benthamiana*, and the fluorescence signals were observed 48 hours-post infiltration using confocal microscopy. Images show the fluorescence from GFP, RFP channel, and the merged fluorescence from both channels. White color in the merged images indicates co-localization. Bars, 20 µm.

To determine whether plant KLCR proteins co-localize with MiEFF17a in plant cells, we examined the subcellular localization of the effector and its host targets. MiEFF17a, without its SP, was fused to Green Fluorescent Protein (GFP) at either its N- or C-terminus, and the constructs were transiently expressed in *N. benthamiana* leaf epidermal cells to identify subcellular compartments of the plant cell targeted by EFF17. When expressed as an N-terminal fusion, GFP-MiEFF17a was mainly detected in the cytoplasm accumulating in bright punctuate structures at the cell periphery (Supplementary Fig. S7). In contrast, the C-terminal GFP fusion displayed a cytoplasmic distribution associated with reticular network reminiscent of the endoplasmic reticulum (ER) (Supplementary Fig. S7). Then, MiEFF17a-GFP was coexpressed with a RFP-ER marker (Nelson *et al*., 2007) or RFP-MiEFF17a together with a MT-Binding Domain (MBD)-GFP marker (Caillaud et al., 2008). These experiments revealed a partial colocalization of MiEFF17a with both ER and MT markers (Supplementary Fig. S7). We subsequently cloned the full length *AtKLCR*s and generated GFP- or RFP-fusions to simultaneously examine the subcellular localization of the effector and its target. Co-expression analysis revealed that MiEFF17a co-localizes with AtKLCR1 predominantly in the cytoplasm and the ER, at the cell periphery (Fig. 3C). Taken together, these findings demonstrate that the EFF17 effector interacts *in planta* with KLCR from both *Arabidopsis* and tomato.

### KLCRs are involved in plant-RKN interaction

Finally, we investigated the potential role of KLCR proteins in RKN parasitism by performing infection assays using *Arabidopsis klcr* mutants. T-DNA insertion mutants have been described for *AtKLCR1* and *AtKLCR2*, previously designated *cmu1* and *cmu2*, respectively, as well as the *cmu1cmu2* double mutant (Ganguly *et al*., 2020). We additionally isolated a new homozygous knock-down line for *AtKLCR3* (SALK_207136; Supplementary Fig. S8), carrying a T-DNA insertion in its promoter, hereafter referred as *cmu3*. The single mutations *cmu1* and *cmu2* did not significantly affect *Arabidopsis* susceptibility to *M. incognita* (Fig. 4), when compared to the Col-0 wild-type. The *cmu3* mutant and *cmu1cmu2* double mutant are less susceptible to the nematode and showed significantly less egg masses (Wilcoxon rank sum test with continuity correction; p-value=0.007103 and p-value=0.003857, respectively) than the wild-type *Arabidopsis* Col-0 (Fig. 4). This result confirms the contribution of KLRs in *Arabidopsis* susceptibility to RKN.

**Fig. 4.**
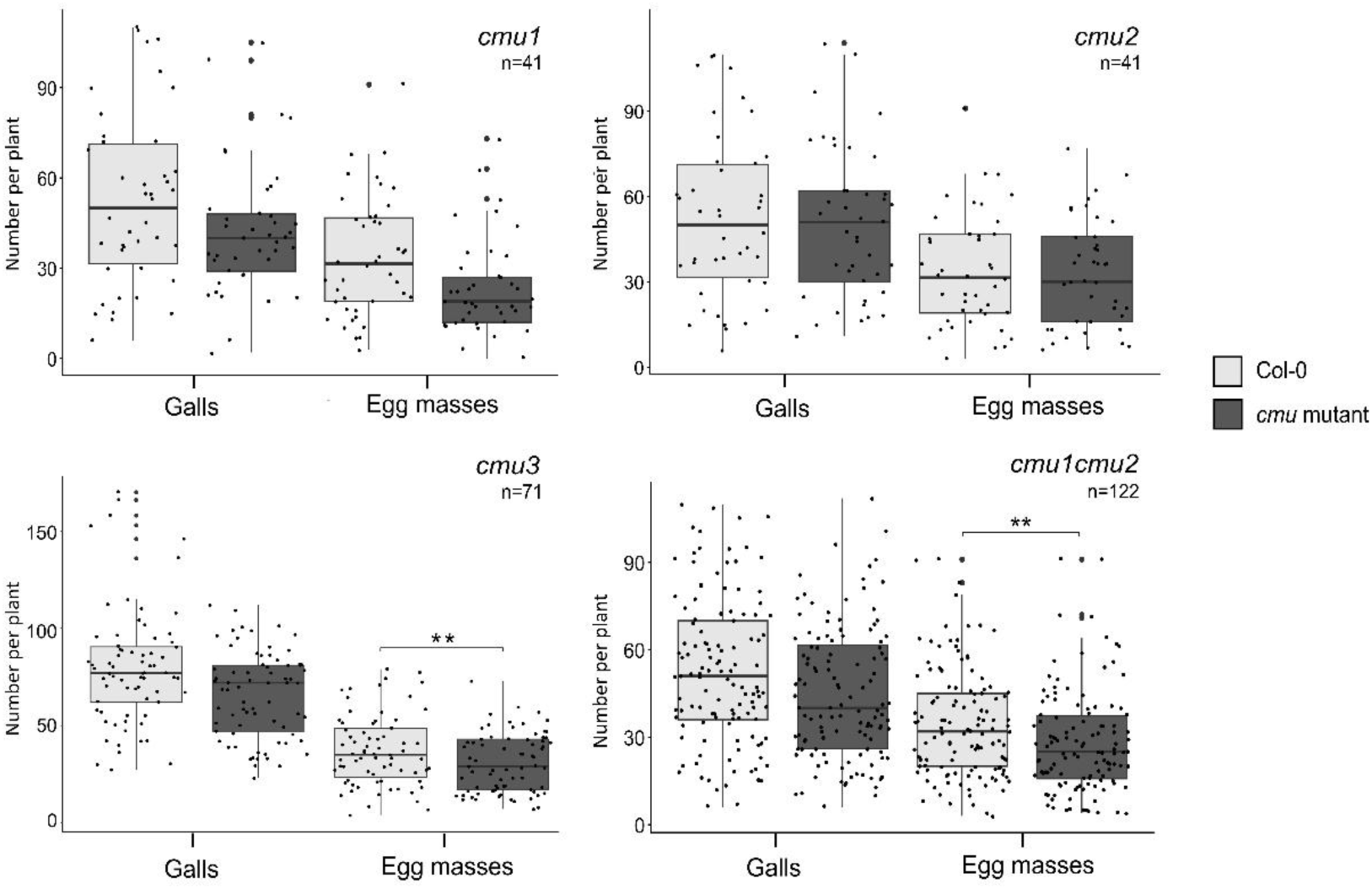
KLCRs contribute to *Arabidopsis* susceptibility to *M. incognita*. Box-and-whisker plots of galls and egg masses per plant in Col-0 control line, *cmu1*, *cmu2, cmu3* and *cmu1cmu2* mutant lines six weeks post infection with 200 *Meloidogyne incognita* second-stage juveniles (J2s). Box indicates interquartile range (25th to the 75th percentile). The central line within the box represents mean value. Whiskers indicate the minimum and maximum values for the normal values present in the dataset. Statistical significance was assessed using Wilcoxon rank sum test with continuity correction (**p<0.01).

## Discussion

RKN of the genus *Meloidogyne* have evolved sophisticated strategies to reprogram host root cells and establish permanent feeding sites. Through the secretion of effector proteins from their esophageal glands, these obligate biotrophs redirect plant developmental programs to induce multinucleate hypertrophied giant cells that sustain their growth and reproduction (Favery *et al*., 2020). The formation and maintenance of these unique feeding structures require extensive remodeling of host cellular architecture, particularly the plant cell wall, the cytoskeleton and endomembrane system (de Almeida Engler *et al*., 2004; Rodiuc *et al*., 2014; Miyashita & Koga, 2017; Orange *et al*., 2025). In this study, we identify EFF17 as a conserved RKN-specific effector that promotes parasitism by targeting plant kinesin light chain-related (KLCR/CMU) proteins, key regulators of cellulose–MT coordination (Liu *et al*., 2016; Ganguly *et al*., 2020).

EFF17 expression in subventral esophageal glands of infective juveniles and its induction during parasitic stages are consistent with a role in host manipulation, as described for other effectors (Vieira & Gleason, 2019; Rutter *et al*., 2022; Bali & Gleason, 2024). Importantly, host-induced gene silencing of MiEFF17 significantly reduced gall formation and egg mass production, demonstrating that this effector contributes to successful parasitism. The conservation of EFF17 across multiple *Meloidogyne* species suggests that it carries out a core function during infection. Conserved effectors are often associated with targeting essential and broadly conserved host processes rather than host-specific adaptations (Mukhtar *et al*., 2011; Weßling *et al*., 2014). Our results support the idea that modulation of cytoskeletal dynamics represents a central and evolutionarily stable strategy used by RKNs to establish feeding sites. Consistently, several nematode effectors have been shown to interfere with cytoskeletal organization, highlighting the importance of structural reprogramming during parasitism (Leelarasamee *et al*., 2018; Mei *et al*., 2018; Godinho Mendes *et al*., 2021).

Plant KLCRs are similar in structure to mammalian Kinesin Light Chains (KLCs) and contain tetratricopeptide repeat (TPR) domains. In plants, KLCRs act primarily as cellulose synthase-MT uncoupling proteins, that stabilize cortical MTs and prevent their lateral displacement caused by the forces exerted by cellulose synthase complexes during cellulose biosynthesis (Liu *et al*., 2016; Ganguly *et al*., 2020). This function is instrumental for maintaining proper cell morphology and overall plant development. In addition, KLCR1 is specifically required for the proper cellulose deposition and architecture in seed mucilage (Yang *et al*., 2022). More broadly, cortical MTs and actin filaments guide the trafficking and spatial targeting of Golgi-derived vesicles carrying matrix polysaccharides to specific sites at the plasma membrane, thereby coordinating their secretion and deposition into the expanding cell wall (Liu *et al*., 2015). AtKLCRs recruit kinesin motors and contribute to these intracellular transport processes (Abel et al., 2013; Ganguly *et al*., 2020). Giant cell formation involves extensive cell expansion and cell wall thickening, conferring the flexibility required for their function as nutrient transfer cells (Rodiuc *et al*., 2014). In healthy roots, cortical MTs guide cellulose microfibril deposition and thus promote anisotropic cell expansion, whereas hypertrophied giant cells expand isotropically (Cabrera *et al*., 2015). Several studies have revealed important changes in cellulose and matrix polysaccharide structure and composition during giant cell ontogenesis (Wieczorek *et al*., 2014; Bozbuga *et al*., 2018; Meidani *et al*., 2019). Manipulation of KLRCs by EFF17 may therefore contribute to reshaping the giant cell wall to favor nematode development.

KLCRs contain TPR domains that mediate protein-protein interactions and promote the assembly of multiprotein complexes (Zeytuni & Zarivach, 2012). Through these complexes, KLCRs participate in linking the cytoskeleton to membranes and integrating cellular signaling pathways. KLCR are key components of membrane contact sites (MCS), particularly the ER-plasma membrane contact sites (EPCS) (Bao *et al*., 2021; Zang *et al*., 2021; Wang *et al*., 2023). KLCR1 interacts with the actin-binding protein NET3C and IQ67-DOMAIN (IQD) proteins to form an actin-MT bridging complex that regulates organelle morphology and dynamics at MCS interface (Zang *et al*., 2021). When plant parasitic nematodes feed by inserting their stylet through the cell wall of host cells, a feeding tube is formed allowing efficient withdrawal of assimilates, which is enclosed by layers of tubular ER (Miyashita *et al*., 2027). Thus, RKNs reshape the host cell’s ER and cytoskeleton to create an organized network that supports their feeding structures. Recent array tomography revealed a potential involvement of MTs in assembling giant cell nuclei into clusters, with ER-rich feeding tubes crossing these nuclear clusters (Orange *et al*., 2025). The cooperation between KLCR and RKN effectors may modulate cytoskeletal and ER organization, thereby facilitating giant cell ontogenesis and/or functionning.

Finally, KLCR proteins are function as central scaffolds in calcium- (Ca2+) mediated cellular signaling. Calmodulins are important Ca2+ sensors that also interact with IQD proteins, which recruit KLCRs to MTs (Abel *et al*., 2013; Bürstenbinder *et al*., 2013; Bao *et al*., 2021; Zang *et al*., 2021). Thus, KLCRs might be crucial contributors to plant responses to biotic triggers and to integrating pathogen perception into reorganization of cellular scaffolds. Consistent with this regulatory role, AtKLCR2 has been identified as a hub targeted by effectors from diverse pathogens, including the bacteria *Pseudomonas synringae, Ralstonia pseudosolanacearum* and *Xanthomonas campestris* and the oomycete *Hyaloperonospora arabidopsidis* (Mukhtar *et al*., 2011; Weßling *et al*., 2014; González-Fuente *et al*., 2020; Supplementary Fig. S9). Moreover, two unrelated RKN effectors, *M. incognita* Minc00344 and *M. javanica* Mj-NULG1a (Supplementary Fig. S10), were shown to interact with the GmHub10, the soybean ortholog of AtKLCR2 (Godinho Mendes *et al*., 2021). These findings underscore the strategic value of host cytoskeletal and membrane trafficking regulators as targets during plant-pathogen interactions (Park *et al*., 2018, Yuen *et al*., 2023).

In conclusion, we propose that EFF17 promotes feeding site ontogenesis by targeting KLCR-dependent scaffolding complexes, thereby reconfiguring cortical MTs and associated membrane domains. Such remodeling may facilitate the isotropic cell expansion and ER reorganization that characterize giant cells. These findings highlight cytoskeletal scaffolding complexes as key susceptibility factors exploited by RKN, and likely other biotrophic pathogens, and provide mechanistic insight into how RKN reshape host cellular architecture to sustain parasitism.

## Materials and Methods

### Nematode and plant materials

*Meloidogyne incognita*, *M. arenaria*, *M. javanica* and *M. enterolobii* were multiplied on *Solanum lycopersicum* cv. “Saint Pierre” and “Piersol”, respectively, grown in a growth chamber (25°C and 16 h photoperiod). Freshly hatched J2s were collected as previously described Caillaud & Favery (2016).

For HIGS experiments, *N. benthamiana* seeds were sown on soil and incubated at 4°C for two days. After germination, plantlets were transplanted into pots containing soil and sand (1:1) and were grown at 24°C, respectively (photoperiod, 16 h: 8 h, light: dark). The *Arabidopsis* lines used for the experiments were from the Columbia (Col-0) genetic backgrounds. The *Arabidopsis* mutant lines *cmu1* (SAIL_335_B08), *cmu2* (SALK_148296), and *cmu1cmu2* double mutant were described by Ganguly *et al*. (2020). The *Arabidopsis* mutant line cmu3 (SALK_207136C) was obtained from the Arabidopsis Biological Resource Center (ABRC, Ohio State University, Columbus, OH, USA). Homozygous mutants were identified by PCR-based genotyping, and new lines were further analyzed by RT-PCR and RT-qPCR, with the primers listed in Supplementary Table 1.

### EFF17 & KLCR sequence analysis, alignment, and phylogenetic tree

The sequences of EFF17 putative paralogues and orthologues were obtained from *Meloidogyne* genomic resources (https://meloidogyne.inrae.fr/) and WormBase ParaSite (https://parasite.wormbase.org/index.html). Sequences of 26 EFF17s belonging to the *M. arenaria*, *M. enterolobii*, *M. floridensis*, *M. graminicola*, *M. incognita*, *M. javanica* and *M. hapla* were identified and aligned using ClustalW algorithm (Thompson *et al*., 2003). Their evolutionary history was inferred by Maximum Likelihood (ML) and the JTT matrix-based model (Jones *et al*., 1992) using MegaX software (Kumar *et al*., 2018). Initial trees calculated for the heuristic search were acquired automatically by applying Neighbor-Join (NJ) and BIONJ algorithms to a pair-wise distance matrix approximated using the JTT model. The evolutionary history was inferred using the Maximum Likelihood method based on the JTT matrix-based model. A discrete Gamma distribution was used to model evolutionary rate differences among sites (5 categories, Γ = 0.68). The bootstrap consensus tree was inferred from 1500 replicates, and branches corresponding to partitions reproduced in less than 50% of bootstrap replicates were collapsed (Felsenstein, 1985). The percentage of replicate trees in which the associated taxa clustered together in the bootstrap test is shown next to the branches. This analysis involved 26 amino acid sequences and a total of 190 positions in the final dataset. Evolutionary analyses were conducted in MEGA11. The pairwise sequence identity matrix of RKN EFF17 protein sequences was generated with Sequence Demarcation Tool version 1.2 software (Muhire *et al*., 2014) (http://web.cbio.uct.ac.za/∼brejnev/). The sequence of *KLCRs* were obtained from The Arabidopsis Information Resource (https://www.arabidopsis.org/) and Sol Genomics Network (https://solgenomics.net/). *KLCR* sequences were aligned using ClustalW algorithm.

### *In situ* hybridization (ISH)

ISH was performed on freshly hatched *M. arenaria*, *M. enterolobii*, *M. incognita* and *M. javanica* preparasitic second stage juveniles, as described previously (Jaouannet *et al*., 2018). For digoxigenin-labelled probes production, the MiEFF17a specific sequences were amplified from entry vectors with the primers MiEFF17_GW3 and MiEFF17_GW5 primers (Supplemental Table S1). An antisense probe was used to detect *EFF17* transcripts in *M. incognita*, *M. arenaria, M. enterolobii* and *M. javanica.* Sense probe was produced that was used as negative control. Images were obtained with a Zeiss Axioplan microscope (Zeiss Axioplan2, Germany).

### Plasmid constructs

The *M. incognita MiEFF17* coding sequences (CDS) lacking the signal peptide and AtKLCR1 were amplified by PCR with specific primers (Supplementary Table S1) and cloned in pDON207 Gateway entry vector (Invitrogen). SlKLCR2, AtKLCR1, AtKLCR2 and AtKLCR3 were synthetized in the entry vector pUC57 with Gateway adapters by Gene Universal Inc (Newark DE, USA). A version of AtKLCR1 with no STOP codon was cloned in pDONR207 vector using specific primers (Table S1). Entry vectors were recombined in the destination vectors pK2GW7 (P35S:MiEFF17a), pK7WGR2 (P35S: mRFP-MiEFF17a), pK7FGW2 (P35S:eGFP-AtKLCR1 and P35S:eGFP-MiEFF17a), pK7FWG2 (P35S:AtKLCR1-eGFP and P35S: MiEFF17a-eGFP), with Gateway technology (Invitrogen). For Split-luciferase assay, the pDEST-Cluc-GWY and pDEST-GWY-Nluc vectors were used (Chen *et al*., 2008). All constructs were sequenced (GATC Biotech) and transferred into *Agrobacterium tumefaciens* strain GV3101.

### Host-induced gene silencing in *Nicotiana benthamiana*

HIGS assays were performed on *N. benthamiana,* as described by Zhao *et al*. (2021). Fragments of MiEFF17a (336 bp) and GFP (298 bp) were amplified by PCR with the primers listed in Supplementary Table S1 and inserted into the TRV2 vector. 10 days post inoculation, plants were inoculated with 200 *M. incognita* juveniles. Six weeks later, roots were stained with 5% eosin and galls and egg masses were counted under a binocular microscope.

### Yeast two-hybrid screening

For yeast two-hybrid (Y2H) screen, the coding sequence (CDS) of the MiEFF17a, without the SP, was used as a bait. MiEFF17a was amplified (Supplementary Table S1) and inserted into the bait vector pB27 (Hybrigenics Service, Paris, France). Y2H screening was performed using an infested tomato root cDNA library (Hybrigenics Service, Paris, France), as described previously (Mejias *et al*., 2021).

### Split luciferase assay

The different N-terminal part of the luciferase (-Nluc) and C-terminal (Cluc-) fusion proteins were transiently expressed in *N. benthamiana* leaves using Agrobacterium-mediated transformation. After 48 hours, leaves were infiltrated with 1mM luciferin (XenoLight D-Luciferin, Perkin Elmer). For quantification, single 4 mm punch disks were placed into wells of a 96 well plate, washed with sterile water and incubated with 1 mM luciferin, as previously (Chen *et al*., 2008). Briefly, after 10 min incubation, light emission was quantified in each well for 5 sec using a luminometer (PerkinElmer, VICTOR Nivo). For each interaction tested, 24 technical replicates were performed (24 leaf disks/plate) and this assay was replicated three times independently. As positive control, Cluc-MSKa and RipG7-Nluc fusion proteins were used as previously described (Wang *et al*., 2016).

### Protein extraction and immunoblotting

*N. benthamiana* leaf discs were ground in liquid nitrogen and proteins were extracted in Laemmli buffer 2X. Yeast protein were extracted after a lysis step in 0.1 M NaOH during 20 min at room temperature followed by protein extraction in protein extraction buffer II (50 mm Tris-HCl pH7.5, 150 mm NaCl, 10 mm EDTA, 0.2% Triton-X-100, 2 mm DTT, 10 mm NaB) and 1 × plant protease inhibitor cocktail (Sigma) and denaturated in Laemmli buffer 2X for 5 min at 95°C. Immunodetection of proteins were performed by loading the samples on precast SDS-PAGE gels (4-15%, Bio-Rad, Hercules, CA, US). Proteins were transferred on nitrocellulose membrane using the Trans-Blot Turbo transfer system (Bio-Rad). The nitrocellulose membranes were blocked with 1X Tris-buffered Saline with Tween20 (TBS-T) solution (137 mM NaCl, 0.1% Tween-20, 3% Milk). Proteins transferred on nitrocellulose membranes were stained in Ponceau S staining solution (0.5% Ponceau S (w/v), 5% acetic acid). Immuno-detection of the protein of interest were performed with the anti-luciferase (1:10000, L0159, Sigma).

### *N. benthamiana* agroinfiltration for *in planta* subcellular localization and split luciferase assay

For cell biology experiments, leaves from 3–4-week-old *N. benthamiana* plants were subjected to agroinfiltration with *A. tumefaciens* recombinant strains as described by Mejias *et al*. (2021). Leaves were imaged 48 hours after agroinfiltration, with an inverted confocal microscope (LSM 880; Zeiss) equipped with a 30 mW argon ion laser and HeNe as the excitation sources. Samples were excited at 488 nm for GFP and 543 nm for RFP. GFP emission was detected selectively with a 505–530 nm band-pass emission filter. We detected RFP fluorescence with a 560–615 nm band-pass emission filter.

For split luciferase assay, OD600=0.125 of each bacterial suspension was used. After incubation at room temperature for 1 h in infiltration medium, bacteria were infiltrated into *N. benthamiana* leaves using a needle-less syringe. The infiltrated plants were incubated in growth chambers under controlled conditions. After 48 h, four leaf disks (8 mm of diameter) were harvested and proteins were extracted for immunoblot analysis.

### RNA Methods

Total RNA was extracted from 5-week-old plants using the RNeasy® Mini kit (Qiagen, Germany), and cDNA synthesis was performed with the SuperScript IV reverse transcriptase (Invitrogen, Thermo Fisher Scientific, Waltham, MA, USA) following the manufacturer’s instructions. For semi-quantitative RT-PCR, cDNA was amplified, with 2 × Rapid Taq Master Mix (Vazyme, China), following manufacturer’s instructions, and the housekeeping *ACTIN2* (*AT3G18780*) was used as an RNA control. For RT-qPCR, PCR reactions were run with the use of the Maxima SYBR Green qPCR Master Mix (29; Fermentas, Thermo Fisher Scientific, Waltham, MA, USA) in an iCycler (Bio-Rad, CA, USA) using gene-specific primers (Supplementary Table S1). *OXA1* (*At5g62050*) and *UBQ10* (*At4g05320*) were used for the normalization of RT-qPCR data. Three technical replicates for three independent biological experiments were performed. Quantifications and statistical analyses were performed with the SATqPCR tool (Rancurel *et al*., 2019)

### Arabidopsis inoculation

Three-week-old *Arabidopsis* plants grown individually in small pots of soil/sand were inoculated with 200 *M. incognita* J2 larvae per plant. Roots were collected 6 weeks after infection, and galls and the egg masses stained with 5% eosin were counted under a binocular microscope. Experiments were done once (*cmu1* and *cmu2*; n=41 plants per line) or repeated three times (*cmu3* and *cmu1cmu2*; n=71 to 122 plants, respectively).

### Graphs and statistical analysis

Graphs and plots were created with R (R.4.4.1 version) and Microsoft® Office Excel® 2019, Jalview (2.11.5.0 version) and GraphPad Prism. Statistical calculations were performed in R (R.4.4.1 version)

## Supporting information

Supplementary informations

## Accession numbers

Sequence data from this article can be found in the The Arabidopsis Information Resource (https://www.arabidopsis.org/), Sol Genomics Network (https://solgenomics.net/) under the following accession numbers: *Meloidogyne incognita MiEFF17a/Minc17612* (*Minc3s01763g26124*), *Arabidopsis thaliana*, *AtKLCR1* (*At4g10840*), *AtKLCR2* (*At3g27960*), *AtKLCR3* (*At1g27500*), *OXA1* (*At5g62050*), *UBQ10* (*At4g05320*), *Solanum lycopersicum SlKLCR1* (*Solyc08g008460*), *SlKLCR2* (*Solyc02g086840*) and *SlKLCR3* (*Solyc03g114320*).

## Acknowledgments

We thank Prof Ram Dixit (University of Saint Louis, Washington, USA) for the kind gift of *cmu1*, c*mu2* and *cmu1cmu2* mutants, and Hybrigenics Services (Paris, France) for providing the pB27 and pP6 vectors and the L40ΔGal4 and Y187 yeast strains. Microscopy work was performed at the SPIBOC imaging facility of Institut Sophia Agrobiotech and we thank Dr Olivier Pierre for his availability.

## Author contributions

SS, NP, BF, MQ conceived and designed the experiments. SS, SM, KM, IM, JP, CC, JM performed the experiments. SS, IM, PA, SJ, NP, BF, MQ analyzed the data. IM, BF, MQ supervised the work. PA, SJ, BF, MQ contributed funding acquisition. SS, BF, MQ wrote the article with input from all authors.

## Data availability

The data that support the findings of this study are available from the corresponding authors upon reasonable request.

## Supplementary material

**Supplementary Table S1.** Specific primers used in this study.

**Supplementary Fig. S1.** Amino acid sequences of EFF17 effector proteins identified in RKN species.

**Supplementary Fig. S2.** MiEFF17 interacts with the three tomato KLCR proteins.

**Supplementary Fig. S3.** Amino acid sequences of tomato KLCR.

**Supplementary Fig. S4.** KLCRs are diversely expressed in tomato tissues.

**Supplementary Fig. S5.** KLCRs are conserved in plants.

**Supplementary Fig. S6.** Western-blot experiments validate the expression of the appropriate Cluc-KLCRs and -Nluc recombinant proteins in the split luciferase assays.

**Supplementary Fig. S7.** MiEFF17a localizes in cytoplasmic structures at the cell periphery and in the ER.

**Supplementary Fig. S8.** Molecular characterization of the *cmu3* T-DNA insertion mutants.

**Supplementary Fig. S9.** AtKLCR2 (AT3G27960) is a hub targeted by several plant pathogen effectors.

**Supplementary Fig. S10.** Plant KLCRs are targeted by three unrelated RKN effectors.

