## Supplementary informations for "A conserved root-knot nematode effector targets plant kinesin light chain related proteins to promote parasitism"

**SUPPORTING INFORMATION**

**Supplementary Table S1.** Specific primers used in this study.

**Supplementary Fig. S1.** Amino acid sequences of EFF17 effector proteins identified in RKN species.

**Supplementary Fig. S2.** MiEFF17 interacts with the three tomato KLCR proteins.

**Supplementary Fig. S3.** Amino acid sequences of tomato KLCR.

**Supplementary Fig. S4.** KLCRs are diversely expressed in tomato tissues.

**Supplementary Fig. S5.** KLCRs are conserved in plants.

**Supplementary Fig. S6.** Western-blot experiments validate the expression of the appropriate Cluc-KLCRs and -Nluc recombinant proteins in the split luciferase assays.

**Supplementary Fig. S7.** MiEFF17a localizes in cytoplasmic structures at the cell periphery and in the ER.

**Supplementary Fig. S8.** Molecular characterization of the cmu3 T-DNA insertion mutants.

**Supplementary Fig. S9.** AtKLCR2 (AT3G27960) is a hub targeted by several plant pathogen effectors.

**Supplementary Fig. S10.** Plant KLCRs are targeted by three unrelated RKN effectors.

**Table S1.** Specific primers used in this study.

| Name | Sequence (5'- 3') | purpose |
|---|---|---|
| MiEFF17_GW5 | AAAAAGCAGGCTTCACCATGTCTGAAATAGGACCAGATAATAAAG | cloning entry vector |
| MiEFF17_GW3_STOP | AGAAAGCTGGGTGTTATCTTCGTCCCCATCCCCAG | cloning entry vector |
| MiEFF17_GW3_noSTOP | AGAAAGCTGGGTGTCTTCGTCCCCATCCCCAG | cloning entry vector |
| AttL1 | TCGCGTTAACGCTAGCATGGATCTC | amplification/sequencing entry vector |
| AttL2 | GTAACATCAGAGATTTTGAGACAC | amplification/sequencing entry vector |
| AttB1 | GGGGACAAGTTTGTACAAAAAAGCAGGCT | amplification/sequencing destination vector |
| AttB2 | GGGGACCACTTTGTACAAGAAAGCTGGGT | amplification/sequencing destination vector |
| MiEFF17-EcoRI-f | CAGAATTCTCTGAAATAGGACCAGATAATAAAG | TRV2 cloning |
| MiEFF17-XhoI-r | GTCTCGAGTCTTCGTCCCCATCCCCAG | TRV2 cloning |
| TRV2-GFP-F | CAGAATTCCATGCCGAGAGTGATCCCG | TRV2 cloning |
| TRV2-GFP-R | GTCTCGAGACGGCAACATCCTGGGGCAC | TRV2 cloning |
| pB27_EFF17_SfiI_NoATG | GGGGCCGGACGGGCCTCTGAAATAGCACCAGATAATAAAG | cloning bait vector |
| pB27_EFF17_SfiI_STOP | AGGGGCCCCAGTGGCCTTATCTTCGTCCCCATCCCCAG | cloning bait vector |
| NP839 | GCCTCCTCTAACGTTCAT | amplification prey vector pP6 |
| NP840 | GCGGGGTTTTTCAGTATC | amplification prey vector pP6 |
| NP841 | TCAAAACCACTGTCACCT | sequencing prey vector pP6 |
| LexA | GTCAGGTCGTTGTCGCACGT | sequencing bait vector pB27 |
| AtKLCR1_GW5 | AAAAAGCAGGCTTCACCATGCCAGCAATGCCAGGTCTCG | cloning entry vector |
| AtKLCR1_GW3_noSTOP | CTTTGTACAAGAAAGCTGGGTGGAACTTGAAACCGAGGCTGGGC | cloning entry vector |
| CLUC | ACGAAGTACCGAAAGGTCTTACC | sequencing split-LUC vector |
| NLUC | TTCATAGCTTCTGCCAACCGAAC | sequencing split-LUC vector |
| LBa1_SALK | TGGTTCACGTAGTGGGCCATCG | genotyping T-DNA insertion line |
| LB1_SAIL | GCCTTTTCAGAAATGGATAAATAGCCTTGCTTGG | genotyping T-DNA insertion line |
| LP_SAIL_335B08_cmu1 | GATCAATCAGATTCTGGAGCG | genotyping T-DNA insertion line |
| RP_SAIL_335B08_cmu1 | TATACAATAAGCCGGTGCCTG | genotyping T-DNA insertion line |
| LP_SALK_148296C_cmu2 | GTTCCTGGAGTCGGTTTTAGG | genotyping T-DNA insertion line |
| RP_SALK_148296C_cmu2 | GTCCAGGGCGAGTTTTATTTC | genotyping T-DNA insertion line |
| LP_SALK_207136_cmu3 | CGACTCCAACGTTCGATTTAG | genotyping T-DNA insertion line |
| RP_SALK_207136_cmu3 | CGTCCATATATCACCATTGGC | genotyping T-DNA insertion line |
| cmu3-f | GAGGAAAAGCTCGGGACAG | RT-PCR |
| cmu3-r | CAAATCTGTGAGAGCCTGTTAA | RT-PCR |
| AtActin2-f | TCAATCATGAAGTGTGATGTGG | RT-PCR |
| AtActin2-r | AACGACCTTAATCTTCATGCTGC | RT-PCR |
| cmu3-f-qPCR | GGGCTTATATGTGAGACTAAGGGAG | RT-qPCR |
| cmu3-r-qPCR | GAACCGAGACAAAGACAAGTACGAG | RT-qPCR |
| OXA1-f | TACCTGATCTGCCTCCACCT | RT-qPCR |
| OXA1-r | GCCAAAGCTGTGGAGAAAAG | RT-qPCR |
| UBQ10-f | CGGAAAACAATTGGAGGATG | RT-qPCR |
| UBQ10-r | TCAAGGGTGATGGTCTTTCC | RT-qPCR |

>MiEFF17a
MNFLNLFLIALIFSFAYASEIGPDNKVKKDEKAEIPTQSKTIEGTVKPEAKPNLVFQSKNKEKSDVAPPFIVKSG
DGKDERQKRWGYGYGGWGGYGGWGGYGGGWGCCGGFGGGWGGGYGYGYGGWGWGRR
>MiEFF17b
MNSIFIKFSLIALILSFAYAAEKGQENEAKKDEKAEILAQSKTIEGAVKPEATPNLVFQSKNKEKSDVAPPFIVK
SGDGKDERQKRWGYGYGGWGGYGGWGGYGGGWGCCGGFGGGWGGGYGYGGWGWGRR
>MiEFF17c
MHFKLQNALLATIRLPTEPLISTFAYAAEKGPENEAKKDEKAEIPAQSKTIEGAVKPEAKPNLVFQSKNKEKSDV
APPFIVKSGDGKDERQKRWGYGYGGWGGYGGYGGWGGYGGGWGCCGGFGGGWGGGYGYGGWGWGRR
>MaEFF17a
MHFKLQNALLATIRLPTEPLILSFAYAAEKGPENEAKKDEKAEIPAQSKTIEGAVKPEAKPNLVFQSKNKEKSDV
APPFIVKSGDGKDERQKRWGYGYGGWGGYGGWGGYGGGWGCCGGFGGGWGGGYGYGGWGWGRR
>MaEFF17b
MHFKLQNALLATIRLPTEPLILSFAYAAEKGPENEAKKDEKAEIPAQSKTIEGAVKPEAKPNLVFQSKNKEKSDV
APPFIVKSGDGKDERQKRWGYGYGGWGGYGGYGGWGGYGGGWGCCGGFGGGWGGGYGYGGWGWGRR
>MaEFF17c
MYFLNLFLIALILSFAYAAEKVPDNEIKKDEKAEIPPHSKTIEGAVKPEAKPNLVFQSKNKEKSDVAPPFIVKSG
DGKDERQKRWGYGYGGWGGGYGGWGGYGGGWGGYGGGWGCCGGGWGGGYGYGYGGWGRR
>MaEFF17d
MNSIFIKFFLIALILSFAYAAEKGPENEAKKDEKAEIPAQSKTIEGAIKPEAKPNLVFQSKNKEKSGVAPPFIVK
SGDGKDERQKRWGYGYGGWGGYGGWGGYGGGWGGGYGGGWGCCGGFGGGWGGGYGYGYGWGRR
>MaEFF17e
NRRGGDISIKYHIFINLSLIALILSFAYAAEKGPNNEVKKDEKAEIPAQSKTIEEAIKPEAKPNLVFQSKNKEKS
DVAPPFIVKSGDGKDERQKRWGYGYGGWGGYGGYGGWGGYGGGWGCCGGFGGGWGGGYGYGGWGWGRR
>MaEFF17f
MHFKLQNALLATIRLPTEPLILSFAYAAEKGQENEAKKDEKAEILAQSKTIEGAVKPEATPNLVFQSKNKEKSDV
APPFIVKSGDGKDERQKRWGYGYGGWGGYGGWGGYGGGWGCCGGFGGGWGGGYGYGGWGWGRR
>MaEFF17g
MHFKLQNALLATIRLPTEPLILSFAYAAEKGQENEAKKDEKAEILAQSKTIEGAVKPEATPNLVFQSKNNEKSDV
APPFIVKSGDGKDERQKRWGYGYGGWGGYGGWGGYGGGWGCCGGFGGGWGGGYGYGYGGWGWGRR
>MjEFF17a
MNFLNLFLIALIFSFAYASEIGPDNKVKKDEKAEIPAQSKTIEGNVKPEAKLNLVFQSKNKEKSDVAPPFIVKSG
DGKDERQKRWGYGYGGWGGYGGWGGYGGGWGCCGGFGGGWGGGYGYGYGGWGWGRR
>MjEFF17b
MHFKLQNALLATIRLPTEPLILSFAYAAEKGPENEAKKDEKAEIPAQSKTIEGAVKPEAKPNLVFQSKNKEKSDV
APPFIVKSGDGKDERQKRWGYGYGGWGGYGGWGGYGGGWGCCGGFGGGWGGGYGYGGWGWGRR
>MjEFF17c
MYFLNLILIALILSFAYAAEKVPDNEIKKDEKAEIPPHSKTIEGTVKPEAKPNLVFQSKNKEKSDVAPPFIVKSG
DGKDERQKRWGYGYGGWGGYGGWGGYGGGWGCCGGFGGGWGGGYGYGGWGWGRR
>MjEFF17d
MHFKLQNALLATIPLILSFAYAAEKGPANEVKKDEKAEIPPQSKTIEGAGKPEAKPNLVFQSKNKEKSDVAPPFI
VKSGDGKDERQKRWGYGYGGWGGYGGWGGYGGGWGCCGGFGGGWGGGYGYGYGGWGWGRR
>MjEFF17e
MNTIFINLSLIALILSFAYAAEKGTENEAKTGEKAEIPAQSKTIDGAVKSEAKPNLVFQSKNKEKSNVAPPFIVK
SGDGKDERQKRWGYGYGGWGGYGGYGGWGGYGGGWGCCGGFGGGWGGGYGYGGWGWGRR
>MfEFF17a
MNTIFINLSLIALILSFAYAAEKGTENEAKTGEKAEIPAQSKTIDGAVKSEAKPNLVFQSKNKEKSDVAPPFIVK
SGDGKDERQKRWGYGYGGWGGYGGYGGWGGYGGGWGCCGGFGWGGYGGGWGCCGGFGGGWGGGYGYGGWGWGRR
>MfEFF17b
MNSIFIKFFLIALISTFAYAAEKGPENEAKKDEKAEIPAQSKTIEGAVKPEAKPNLVFQSKNKEKSDVAPPFIVK
SGDGKDERQKRWGYGY
>MeEFF17a
MNSIFIKFFLIALILSFAYAAEKGPVNEAKKDEKAEITAQSKTIEGAVKPEANPNLIFQSKNKEKSDVAPPFIVK
SGDGKDERQKRWGYGYGGWGGYGGYGVWGGYGGGWGCCGGFGGGWGGGYGYGGWGWGRR
>MeEFF17b
MNSIFIKFSLIALILSFAYAAEKGQENEAKKDEKAEILAQSKTIEGAVKPEATPNLVFQSKNKEKSDVAPPFIVK
SGDGKDERQKRWGYGYGGWGGYGGWGGYGGGWGCCGGFGGGWGGGYGYGGWGWGRR
>MeEFF17c
MNSIFIKFFLIALILSFAYAAEKGPVNEAKKDEKAEILAQSKTIEGAVKPEATPNLVFQSKNKEKSDVAPPFIVK
SGDGKDERQKRWGYGYGGWGGYGGWGGYGGGWGCCGGFGGGWGGGYGYGGWGWGRR
>MeEFF17d

```
MYFLNIFLIALILSFAYAAEKVPDNEIKKDEKAEIPPHSKTIEGTVKPEAKPNLVFQSKNKEKSDVAPPFIVKSG
DGKDERQKRWGYGYGGWGGYGGWGGYGGGWGCCGGFGGGWGGGYGYGGWGWGRR
>MeEFF17e
MNSIFIKFFLIALILSFAYAAEKGPDNEVKKDEKAEIPAQSKTIEGAIKPEAKPNLVFQSKNKEKSDVAPPFIVK
SGDGKDERQKRWGYGYGGWGGYGGWGGYGGGWGWGGGYGGGWGCCGGYGYGGWGWGRR
>MeEFF17f
MNSIFIKSFLIALLLSFAYAAEKGPDNEVKKDEKAEIPDQTKTIEGAVKPEAKPNLVFQSKNKEKSDVAPPFIVK
SGDGKDERQKRWGYGYGGWGGYGGWGGFGGGWGWGGGYGGGWGCCGGYGYGGWGWGRR
>MeEFF17g
MNFLNLFLIALIFSFAYASEIGPDNKVKKDEKAEIPTQSKTIEGTVKPEAKPNLVFQSKNKEKSDVAPPFIVKSG
DGKDERQKRWGYGYGGWGGYGGWGGYGGGWDVVEALVEDGWRIWLWLWWLGMGTKIKSIVKIT
>MhEFF17a
MNNIFAEKGPNNEVKKDEKAEIPAQSKTIEEAIKPEAKPNLVFQSKNKEKSDVAPPFIVKSGDGKDERQKRWGYG
YGGWGGYGGYGVGAVMEVDGDVVEALVEDGVADMVMVAGDGDEDKKHCQNNIGVNLNELTTE
>MgEFF17a
MKILLTIFFISLILAFTYAKEKIPENEGEKDEKTLLDAKKEENPINPEIKPNLVFQAKAPAKAADGPVFTEKTGD
GKEGRHKRWWGGWGGCCGGWGGWGGYGGWGGWGGYGGYGGYGWGWGRR
```

**Supplementary Figure S1.** Amino acid sequences of EFF17 effector proteins identified in RKN species.

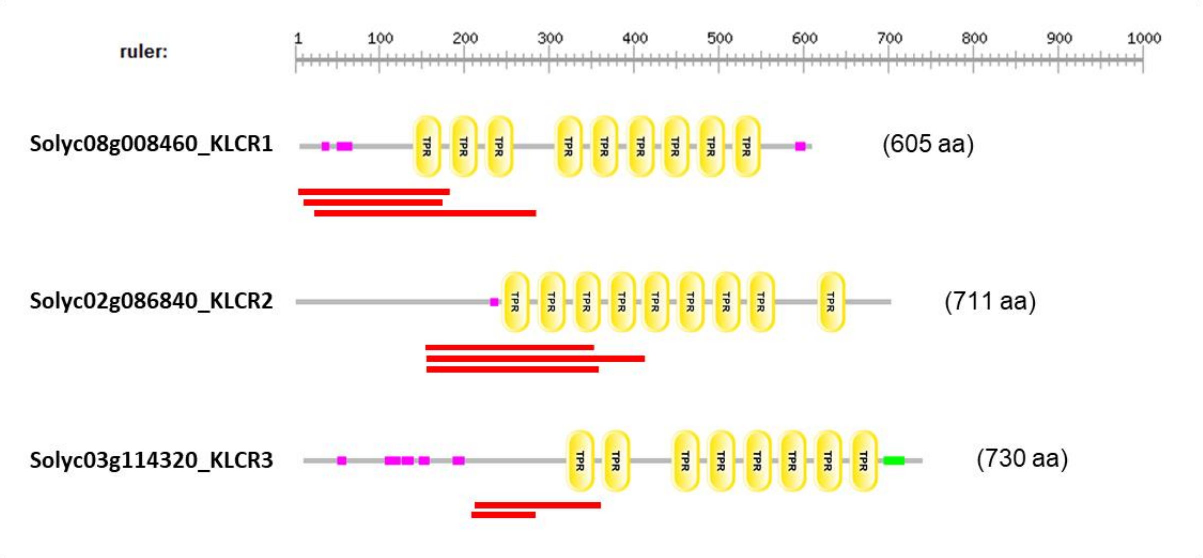


**Supplementary Figure S2.** MiEFF17 interacts with the three tomato KLCR proteins. The schematic representation of the three tomato KLCR proteins was obtained using the Simple Modular Architecture Research Tool (SMART) (http://smart.embl-heidelberg.de/), and reveals presence of conserved tetratricopeptide repeat (TRP) domains (in yellow; IPR019734). The different peptides captured in the yeast-two hybrids screening are displayed as red lines. In pink, low complexity regions. In green, coiled coil region.

```
>SOLYC08G008460/SlKLCR1
MPGLVSVKTPPETPALRISISDENHGRNGSGSSRSEQFNPKTNSPAPRRPPSPSTSRAKPSPDRGSGKKKSPPEK
VEIEESFLDNPDLGPFLLKLARDTIASGEGPTKALDYALRAAKSFERCAVDGEPSLDLAMSLHVVAAIYCSLGRF
DEAIPVLETAIKVPEVSRGADHALAAFSGYMQLGDTYSMLGQLDRSIESYKEGLKTQMEALGDTDPRVAETCRYL
AEAHVQAMQFDEAENLCKKTLEIHRAHSPPASLEEAADRRLMALICEAKCDYESALEHLVLANMAMIANGQETEV
AAIDVGIGNIYLSLSRFDEAVFSYQKALTVFKSSKGDNHPSVASVYVRLADLYYKTGKLRESRSYCENALRIYAK
PVPGTTPEDIACGLTEISAVYELFNEPEEALKLLLKAMKLLEDKPGQQSTIAGIEARMGVMFYMVGRYEEARSSL
ENAVIKLRASGERKSAFFGVVLNQMGLSSVQLFKIDEAAELFEEAKEILEQECGHCHQDTLGVYSNLAATYDAIG
RVDDAIEILEYVLKLREEKLGTANPDFNDEKKRLAELLKEAGRSRNKNPKSLENLIDPNSKRTTKKETSSKKWSA
FGFRS
>SOLYC02G086840/SlKLCR2
MDGSIVDGNQKEPNGHFLPHREIFDQGSPRSPLSTHSRETESIDLDINGGVDTSIEQLYNNVYEMQSSDYSPSRR
SFLSYGEESRIDSELRYLAGGDFGELDSKKGLSEHDKVHNDEKLGKIKTYPASPKSVWSAKGKKYSPSRTDSPIA
NKPPRSRSKSFNEKPSPKRFGNLKKLNATMSMKNEKNPNANEDSSKAGYLGPYLLKQARDMISTTGENVQKALEL
ALRAMKSFESSSKGNSSLEFVMCLHVVAALHCRLGKYNEAIPLLERSIEIPDLDVGQNHALAKFAGCMQLGDTYA
MLGQLENSILCYTAGLEIQRQVLGEKDTRFGETCRYVAEAHVQAMQFDEAEKLCQMALDIHKENNSSASPEEAAD
RRLLGLIYDSKGDYEAALEHYVLAGMAMAANGQEADVASIDCNIGDAYLSMARYDEAICAYQKALTKFKSTKGEN
HPSVASVYVRLADLYNKIGKFRESKSYCENALRIYTKAVPGSHPEEIASGLVDVSVIYESMNEPDQALKLLQKAI
KVYGNAPGQQSTIAGIEAQIGVLYYILGDYMDSYDSLKTAVSKFREIGEKKSAIFGITLNQMGLACVQLYAINEA
GDLFEEARIILETECGPYHADTLGIYSNLAGTYDAMGRTDDAIEILEFVVGMREEKLGTANPDVDDEKRRLTELL
RESGRVRSRKSRSLETLLGNMSHIFLQDQEIDILER
>SOLYC03G114320/SlKLCR3
MPGVVMDEIHEVGEVKELKENGNSTPCKENEEGGLGPRNGGEEHVGDGVVEPSIEELYENVCEMQSSDQSPSRHS
FGSDGDESRIDSELRHLVGGEMREVEIIEEDEEVQKPEIEDSRSDSGSKKGTSDDVKLDNSPSSSTKDPSSGQPK
TPSQLELESETSAKSNSKGRRASLDKKNGNNTKKVVVGGTSRSRQKSSPASGSKLKNGTEDSSDSGLDNPDLGPF
LLKQARDLIASGDNSHKALELAHRAAKSFEKCANGKPSLDVVMCLHVTAAIYCNLGQYDDAIPLIEHSLEIPVVE
EGQEHALAKFAGYMQLGDTYAMLGQLENSIVSYTTGMEIQRQVLGDSDPRVGETCRYLAEAHVQALQFDEAEKLC
QMALDIHKENGSPPSLEEAADRRLMGLICESKGDHEAALEHLVLASMAMVANGQESEVASVDCSIGDTYLSLNRY
DEAIFAYQKALTALKSSKGENHPAVASVFVRLADLYNRTGKLRDSKSYCENALRIYGKPIPGIAPEEIANGLTDV
SAIYESMNELDQALKLLQRALKIYNNAPGQQNTIAGIEAQMGVIYYMLGKYSESYNSFKSAISKLRASGEKKSAF
FGVALNQMGLACVQRYAINEAVELFEESKVILEQEYGPYHPETLGVYSNLAGTYDAVGRLDEAIEILEYIVGVRE
EKLGTANPDVADEKKRLAELLKEAGRVRNRKARSLENLLDANHRPNAINNDLIIV
```

**Supplementary Figure S3.** Amino acid sequences of tomato KLCR.

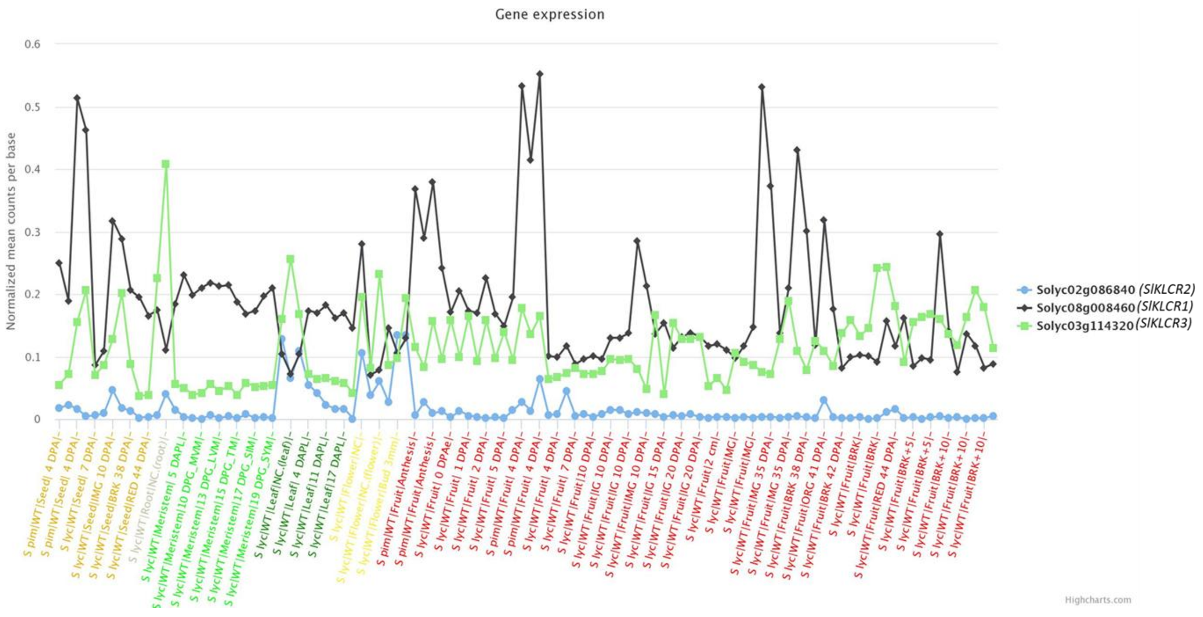


**Supplementary Figure S4.** *KLCRs* are diversely expressed in tomato tissues. Expression data were retrieved from TomExpress RNA-seq platform. The expression of each *SlKLCR* gene is shown in wild type (WT) *S. lycopersicum* (*S lyc*) and *S. pimpinellifolium* (*S pim*). Gene transcription was analyzed in seeds, roots, leaves, meristems (middle vegetative meristems, MVM; late vegetative meristems, LVM; transition meristems, TM; sympodial inflorescence meristems, SIM; sympodial shoot meristems, SYM;), flowers and fruits (immature green, IG; mature green, MG; breaker, BRK; orange, ORG; red ripe, RR). DPA, days post anthesis; DPAe, 0 DPA; DPG, days post germination; DAPL, days after planting; NC, not communicated; the number after "+" indicate number of days. Additional information on normalization methods, the tissue, and the developmental stages of each sample can be found at https://tomexpress.gbfwebtools.fr/docs.

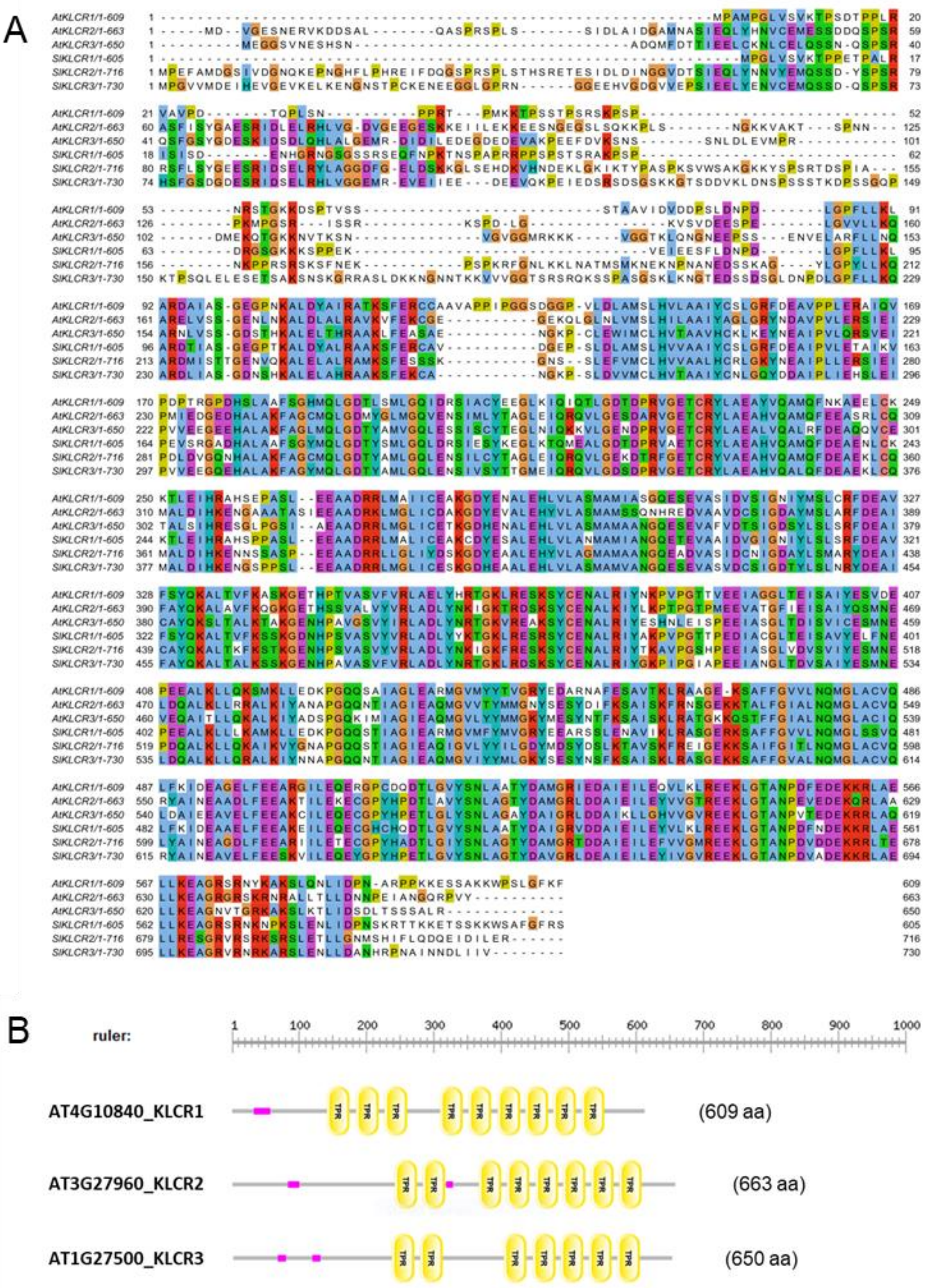


**Supplementary Figure S5.** KLCRs are conserved in plants. **A,** Alignment of full-length sequences of tomato and *Arabidopsis* KLCR proteins created using ClustalX and visualized using Jalview with the ClustalX default colors. **B,** Schematic representation of the three *Arabidopsis* KLCR proteins was obtained using the Simple Modular Architecture Research Tool (SMART) (http://smart.embl-heidelberg.de/) and reveals presence of conserved tetratricopeptide repeat (TRP) domains (in yellow; IPR019734). In pink, low complexity regions.

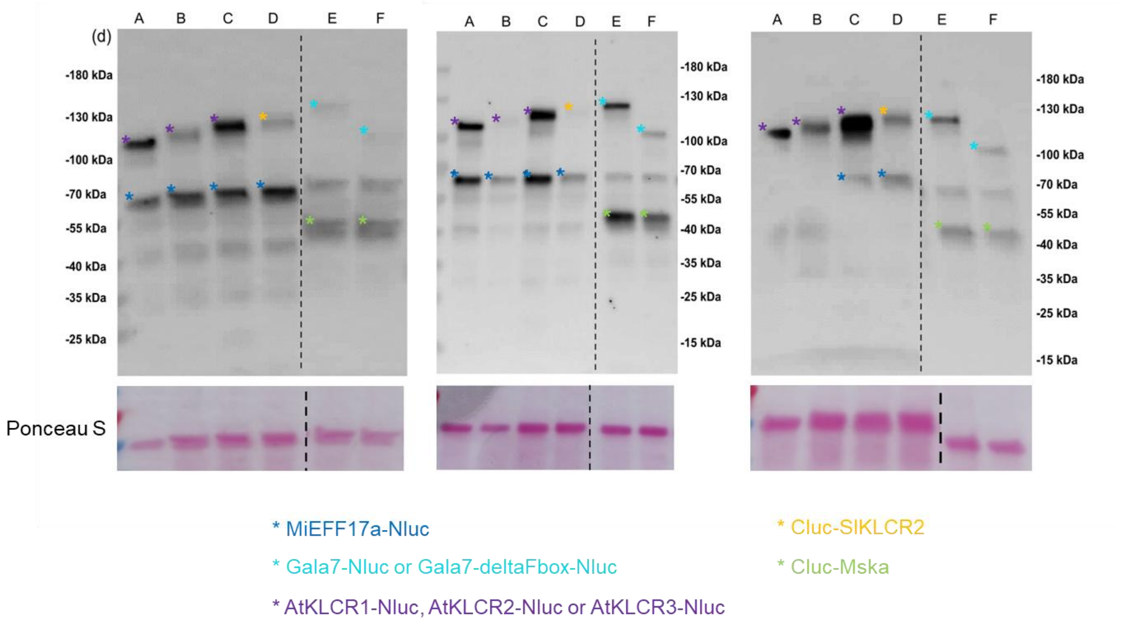


**Supplementary Figure S6.** Western-blot experiments validate the expression of the appropriate Cluc-KLCRs and -Nluc recombinant proteins in the split luciferase assays (related to Figure 3). 48 hours-post agro-infiltration of *N. benthamiana* leaves, Cluc- and -Nluc recombinant proteins were extracted and analyzed by immunoblotting with anti-luciferase antibodies to detect Cluc and Nluc. Results from three independent experiments show similar results. A, MiEFF17a-Nluc + Cluc-AtKLCR1; B, MiEFF17a-Nluc + Cluc-AtKLCR2; C, MiEFF17a-Nluc + Cluc-AtKLCR3; D, MiEFF17a-Nluc + Cluc-SlKLCR2; E, Gala7-Nluc + Cluc-MSKa; Gala7-deltaFbox-Nluc + Cluc-MSKa.

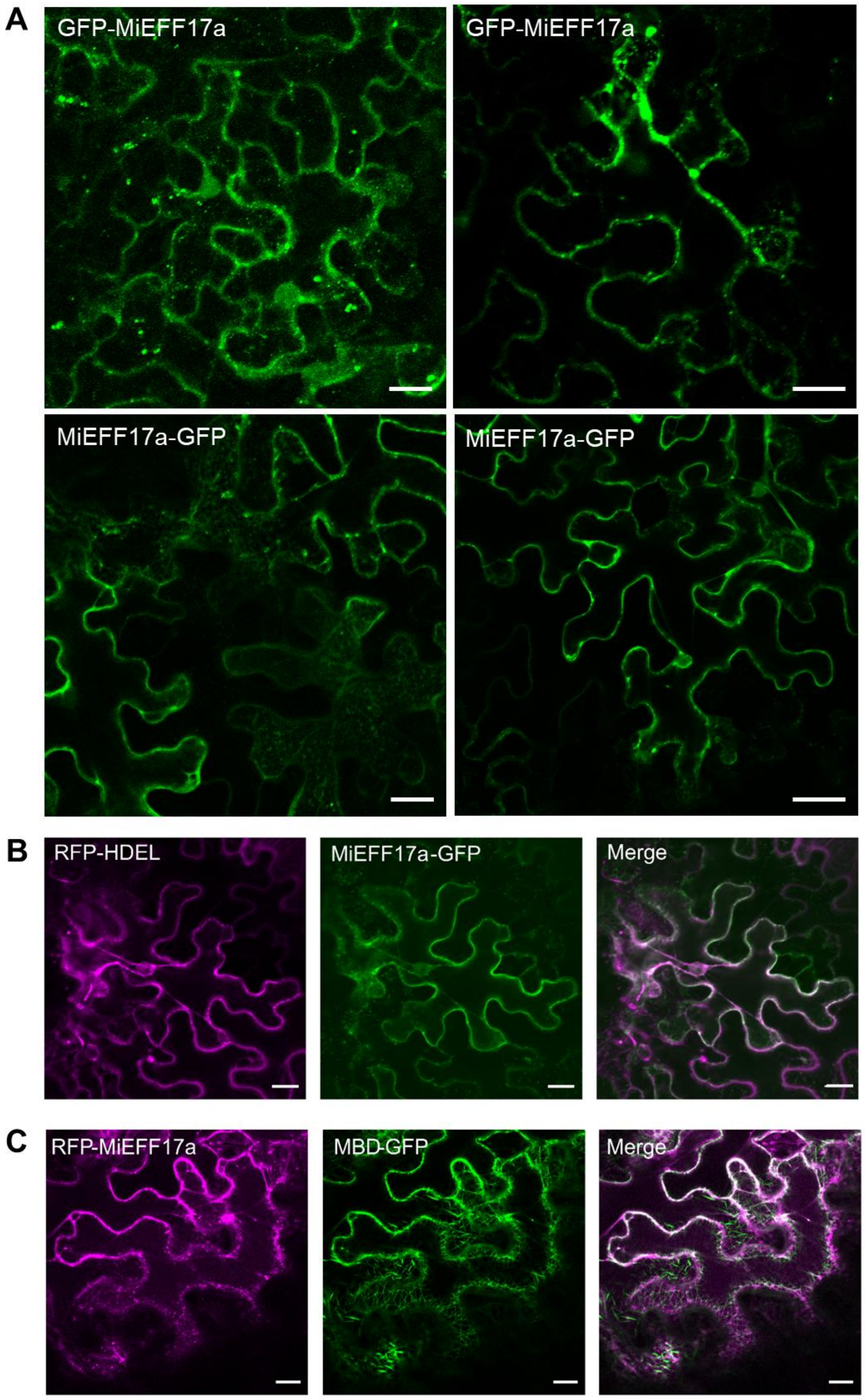


**Supplementary Figure S7.** MiEFF17a localizes in cytoplasmic structures at the cell periphery and in the ER. **A,** Laser-scanning confocal micrographs showing localization of GFP-MiEFF17a and MiEFF17a-GFP recombinant protein transiently produced in N. benthamiana agro-infiltrated leaves. **B-C,** Co-localization analysis of MiEFF17a with an ER (HDEL) or a microtubule (MBD) marker. MiEFF17a-GFP and RFP-HDEL (**B**) or RFP-MiEFF17a and MBD-GFP (**C**) were transiently co-expressed using *Agrobacterium* in *N. benthamiana*, and the fluorescence signals were observed 48 hours-post infiltration using confocal microscopy. Images show the fluorescence from GFP, RFP channel, and the merged fluorescence from both channels. White color in the merged images indicates co-localization. Bars, 20 µm.

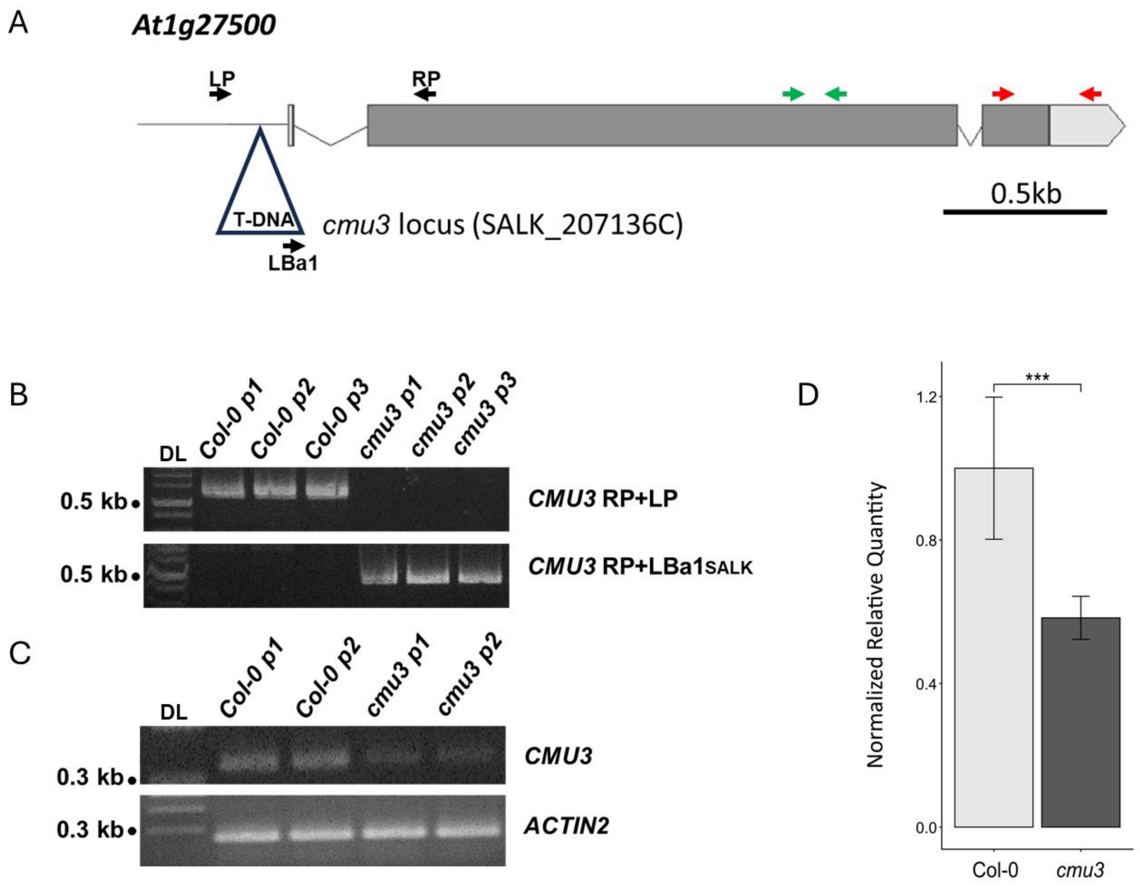


**Supplementary Figure S8.** Molecular characterization of the *cmu3* T-DNA insertion mutants. **A,** Schematic representation of the *cmu3* locus carrying a T-DNA insertion in the promoter of the *At1g27500* gene. The position of primers used for genotyping (LP, RP and LBa1), for RT-PCR (red arrows) and RT-qPCR (green arrows) is displayed. **B,** PCR on genomic DNA from three randomly selected wild-type Arabidopsis plants (Col-0) and three *cmu3* T-DNA insertion mutant (p1 to p3) validates the insertion of the T-DNA in the promoter of the *At1g27500* gene in the *cmu3* homozygous line. DL, DNA ladder. **C,** *CMU3* transcripts in two randomly selected Col-0 and *cmu3* mutant plants (p1 and p2) were quantified by RT-PCR analysis. *ACTIN2* was used as an RNA control. DL, DNA ladder. **D,** *CMU3* transcripts in Col-0 and *cmu3* line were quantified by RTqPCR. Shown are normalized relative transcript quantities calculated with the SatqPCR software from three independent biological replicates. The *AtOXA1* and *AtUBQ10* housekeeping genes were used for data normalization. Error bars indicate the SD.

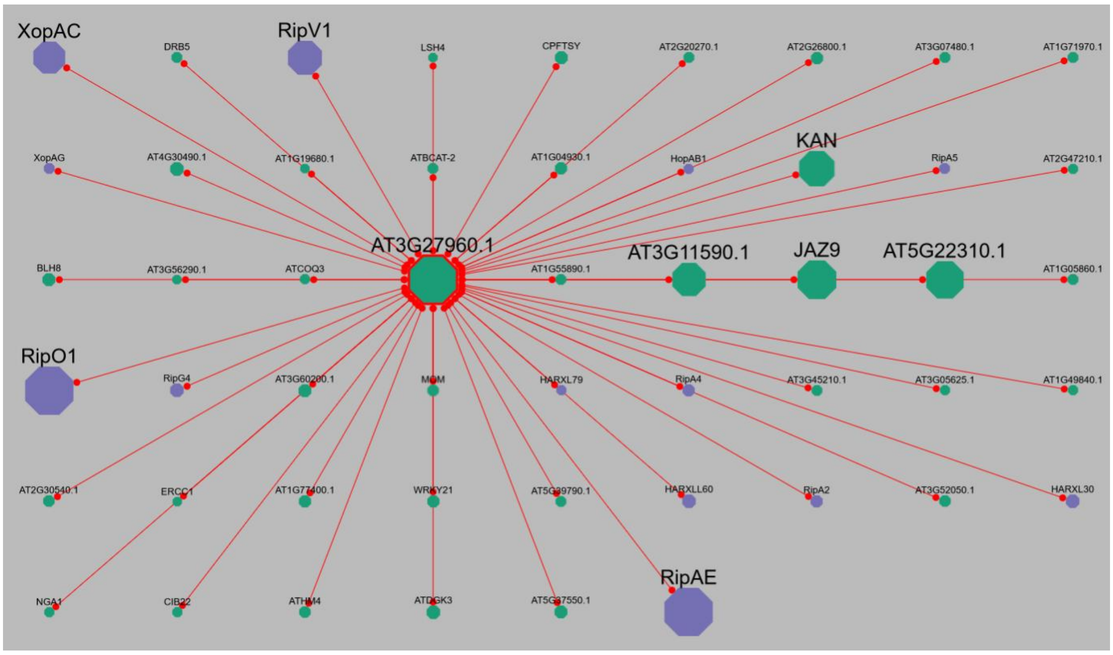


**Supplementary Figure S9.** AtKLCR2 (AT3G27960) is a hub targeted by several plant pathogen effectors. In purple, effectors form the bacteria *Xanthomonas campestris* (XopAC and XopAG), *Ralstonia pseudosolanacearum* (RipV1, RipA5, RipO1, RipG4, RipA4, RipA2 and RipAE) and *Pseudomonas synringae* (HopAB1), and the oomycete *Hyaloperonospora arabidopsidis* (HARXL79, HARXLL60 and HARXL30). In green, interactors in *Arabidopsis thaliana*. Data were collected using EffectorK (https://lipm-browsers.toulouse.inrae.fr/k/EffectorK/; González-Fuente *et al.*, 2020).

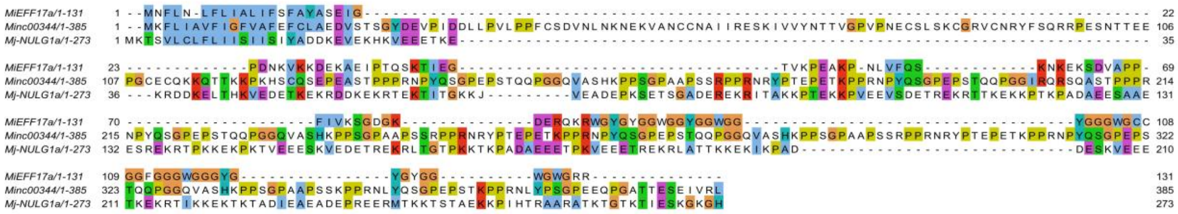

**Supplementary Figure S10.** Plant KLCRs are targeted by three unrelated RKN effectors. MiEFF17a amino acid sequence was aligned with sequences of Minc00344 and Mj-NULG1a effectors also known to interact with plant KLCRs (Godinho Mendes *et al.*, 2021).
